# GetPrimers: a generalized PCR-based genetic targeting primer designer enabling easy and standardized targeted gene modification across multiple systems

**DOI:** 10.1101/2023.08.12.552711

**Authors:** Zepu Miao, Haiting Wang, Xinyu Tu, Zhengshen Huang, Shujing Huang, Xinxin Zhang, Fan Wang, Zhishen Huang, Huihui Li, Yue Jiao, Song Gao, Zhipeng Zhou, Chun-Min Shan, Jing Li, Jia-Xing Yue

## Abstract

Genetic targeting (e.g., gene knockout and tagging) based on polymerase chain reaction (PCR) is a simple yet powerful approach for studying gene functions. Although originally developed in classic budding and fission yeast models, the same principle applies to other eukaryotic systems with efficient homologous recombination. One-step-PCR-based genetic targeting is conventionally used but the sizes of the homologous arms that it generates for recombination-mediated genetic targeting are usually limited. Alternatively, gene targeting can also be performed via fusion PCR, which can create homologous arms that are orders of magnitude larger, therefore substantially increases the efficiency of recombination-mediated genetic targeting. Here we present GetPrimers (https://www.evomicslab.org/app/getprimers/), a generalized computational framework and web tool to assist automatic targeting and verification primer design for both one-step-PCR-based and fusion-PCR-based genetic targeting experiments. Moreover, GetPrimers by design runs for any given genetic background of any species with full genome scalability. Therefore, GetPrimers is capable of empowering high-throughput functional genomics assays at multi-population and multi-species levels. Comprehensive experimental validations have been performed for targeting and verification primers designed by GetPrimers across multiple organism systems and experimental setups. We anticipate GetPrimers to become a highly useful and popular tool to facilitate easy and standardized gene modification across multiple systems.

**Take Aways:** Developed GetPrimers for generalized primer design of PCR-based genetic targeting.

GetPrimers shines with its high versatility and genome-wide scalability.

GetPrimers was validated across multiple organisms and experimental setups.

## Introduction

The long-standing promise of genetics is to establish the map between genotypes and phenotypes. This was traditionally done by dissecting and mapping the genetic basis of observed phenotypes, a process known as “forward genetics”. In the era of functional genomics, when genome sequences become readily available, researchers can take the “reverse genetics” approach by directly modifying, deleting, or tagging a specific gene locus and examining the corresponding phenotypic changes. Most gene-targeting strategies in fungi are based on the one-step gene disruption technique developed first in *S. cerevisiae* [Rothstein 1983]. Such precise gene targeting is often accomplished by swapping in a targeting cassette to replace the original genomic fragment through homologous recombination. For the targeting cassette, heterologous antibiotic selection markers such as *kanMX* or *hphMX* are often used to select for successful integration. In the case of gene tagging, extra sequences coding for fluorescent proteins or epitope tags are also needed as a part of the targeting cassette, with which homologous recombination will generate in-frame fusion protein with the targeted gene during translation. For N-terminal tagging, a regulatable promoter is usually added into the cassette as well to fine-control the expression level of the tagged gene. Nowadays, homologous recombination-mediated gene targeting has become a routine experiment for labs world-wide working on different organisms.

For organisms with highly efficient homologous recombination machinery (e.g., the budding and fission yeasts), gene targeting experiments can be conveniently performed with one-step Polymerase chain reaction (PCR), during which target-matching homologous arms are introduced with the designed PCR primers. Such introduced homologous arms will guide the precise integration of the construct that carries the gene deletion or tagging cassette. For the budding yeast *Saccharomyces cerevisiae*, PCR-added homologous arms of 35-40 bp on both sides of the target locus are enough to trigger efficient sequence integration [Baudin, Ozier-Kalogeropoulos, Denouel, Lacroute, and Cullin 1993; Wach, Brachat, Pöhlmann, and Philippsen 1994; Goldstein and McCusker 1999]. The systematic application of this clone-free technique at the whole genome level empowered the landmark functional genomics studies such as the Yeast Deletion Collection Project [Giaever et al. 2002; Winzeler et al. 1999]. For the fission yeast *Schizosaccharomyces pombe*, longer homologous arms (e.g., 80-100 bp) are usually recommended given its comparatively lower homologous recombination efficiency [Bähler, Wu, Longtine, Shah, Mckenzie III, Steever, Wach, Philippsen, and Pringle 1998]. Similar strategies have also been applied to other yeast and filamentous fungi species, such as *Kluyveromyces lactis* [Kooistra, Hooykaas, and Steensma 2004], *Cryptococcus neoformans*[Goins, Gerik, and Lodge 2006], *Ashbya gossypii* [Wendland, Ayad-Durieux, Knechtle, Rebischung, and Philippsen 2000], and *Neurospora crassa* [Colot, Park, Turner, Ringelberg, Crew, Litvinkova, Weiss, Borkovich, and Dunlap 2006], although much longer homologous arms (e.g., 1-kb long for *Neurospora crassa*) are sometimes needed to ensure a satisfiable integration rate. Synthesizing long-arm-carrying PCR primers can be quite costly and PCR experiments with such very long primers can also be tricky. These practical challenges motivated the invention of fusion-PCR-based gene targeting methods [Amberg, Botstein, and Beasley 1995; Nikawa and Kawabata 1998; Kuwayama, Obara, Morio, Katoh, Urushihara, and Tanaka 2002], for which researchers first used PCR reactions to amplify the two homologous arms and then let them fuse with the central PCR fragments that carries the gene targeting cassette. Such fusion-PCR-based gene targeting strategy substantially broaden the application scope of PCR-based gene targeting methods. Successful application of this method has been reported for many fungi organisms with less efficient homologous recombination machineries [You, Lee, and Chung 2009; Fu, Hettler, and Wickes 2006] as well as for non-fungi species such as the social amoeba *Dictyostelium discoideum* [Kuwayama et al. 2002].

Despite differences in their technical details, one-step-PCR- and fusion-PCR-based gene targeting methods share one prerequisite in common, which is the meticulous design of targeting and validation PCR primers. In many labs, this is typically done manually. For one-step-PCR-based gene targeting, this is relatively straightforward given that the gene targeting PCR primers are co-determined by gene flanking sequences and common plasmid primers. Yet it could be laborious when the number of genes to be targeted scales up. Moreover, to verify whether the cassette integration is correct and precise, researchers still must manually design verification primers for each targeted gene locus. Therefore, a couple of bioinformatic tools have been developed to perform automatic primer designing based on the reference genome of *Saccharomyces cerevisiae* and *Schizosaccharomyces pombe* [Penkett, Birtle, and Bähler 2006; Yofe and Schuldiner 2014; Cummings, Joh, and Motamedi 2015; Wang, Xu, Wang, Liu, Lou, and Sugiyama 2021]. Unfortunately, for fusion-PCR-based gene targeting method, no automatic primer design tool is available so far. Given that the primer design of fusion-PCR-based gene targeting is substantially more complicated and error-prone, such lack of automated primer design tools really limited the application of fusion-PCR-based gene targeting, which is more versatile and powerful.

Here we developed GetPrimers (https://www.evomicslab.org/app/getprimers/), a generalized computational framework and web tool to perform automatic targeting and verification primer design for PCR-based gene targeting. Serials of generalization have been applied during our method design and software implementation. 1) GetPrimers supports gene targeting based on both one-step PCR (i.e., the “long primer” strategy) and fusion PCR (i.e., the “short primer” strategy), enabling users to freely explore both methods based on their specific use scenarios. 2) GetPrimers supports locus-specific, batch-specific, and genome-wide primer design, empowering large-scale experimental design with full scalability. 3) While gene targeting is mainly for targeting protein-coding genes as its name implies, GetPrimers can also design primers for targeting any user defined genomic locus. 4) GetPrimers runs on any given species and any given genetic background, facilitating studying gene function across expanded genomic and phenotypic spaces at both population and species levels. Taken together, we anticipate GetPrimers to become a highly capable and useful tool to facilitate researchers using PCR-based gene targeting in yeasts, filamentous fungi, and other suitable eukaryotic species.

## Materials and Methods

### Software implementation and web server deployment

The GetPrimers command-line program was written in Perl and hosted at GitHub (https://github.com/codeatcg/GetPrimers). Third-party tools such as ncbi-blast+[Camacho, Coulouris, Avagyan, Ma, Papadopoulos, Bealer, and Madden 2009] (v2.11.0), WindowMasker[Morgulis, Gertz, Schäffer, and Agarwala 2006] (v2.11.0), and Primer3[Untergasser, Cutcutache, Koressaar, Ye, Faircloth, Remm, and Rozen 2012] (v2.6.1) were used to facilitate GetPrimers to perform its task. To make the program easier to use, a dedicated web interface was created on top of the command-line program by using HTML, jQuery and Bootstrap. Django was used as backend and an SQLite3 database was created for storing pre-calculated primer combinations. The GetPrimers webserver is deployed via an Alibaba simple application server with 2 CPU and 4 GB RAM, which is globally accessible with the URL: https://www.evomicslab.org/app/getprimers/.

For the webserver, genome-wide targeting and verification primer combinations were pre-calculated for representative strains of *Saccharomyces cerevisiae*, *Saccharomyces paradoxus*, *Schizosaccharomyces pombe*, *Neurospora crassa* and a few other fungi species (including multiple plant pathogens) (Table 1). The genome assembly and annotation of more species and strains will be continuously added upon users’ request.

**Table 1.**
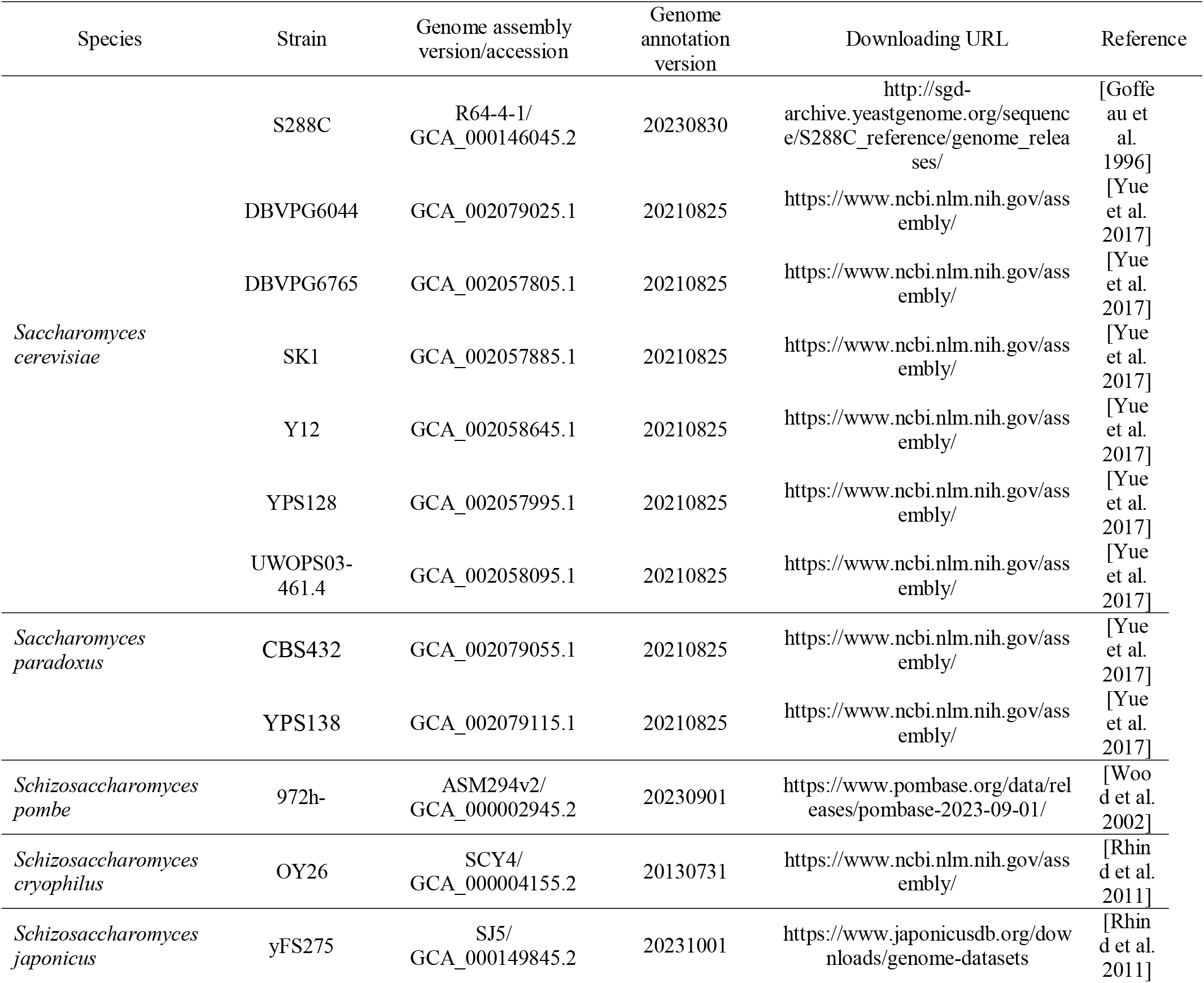
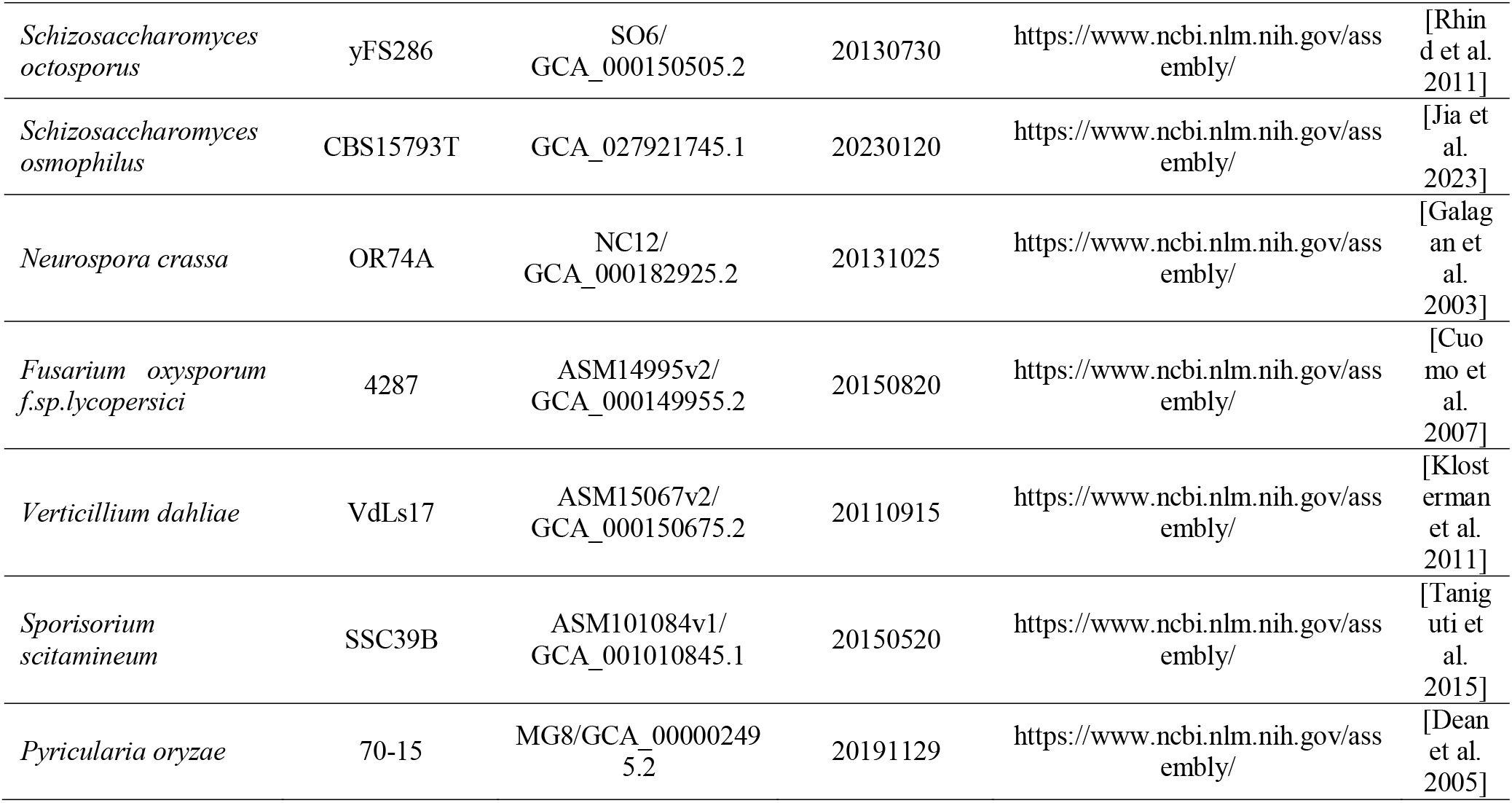
Pre-shipped genome assemblies used for GetPrimers’ webserver.

### Strains and medium

All species and strains used in this study and their respective ploidy and genotype information are provided in Table 2.

**Table 2.** The species and strains used in the application demonstration for GetPrimers.

| Species | Strain | Ploidy | Genotype |
| --- | --- | --- | --- |
| *Saccharomyces cerevisiae* | BY4741 | 1n | MATa, his3Δ1, leu2Δ0, lys2Δ0, ura3Δ0 |
|  | DBVPG6044 | 1n | MATa, his3Δ1, leu2Δ0, lys2Δ0, ura3Δ0 |
|  | Y12 | 1n | MATa, ura3Δ0, leu2Δ0, lys2Δ0, met15Δ0 |
|  | YPS128 | 1n | MATa, ura3Δ0, leu2Δ0, lys2Δ0, met15Δ0 |
|  | UWOPS03-461.4 | 1n | MATa, ura3Δ0, leu2Δ0, lys2Δ0, met15Δ0 |
| *Saccharomyces paradoxus* | CBS432 | 2n | Wildtype |
|  | YPS138 | 2n | Wildtype |
| *Schizosaccharomyces pombe* | SPJ649 | 1n | h- smt0, leu1-32, his2, ura4DS/E, ade6-M210 |
| *Neurospora crassa* | *Ku70<sup>RIP</sup>* | 1n | *bd, a* |

The culture medium used in this study is listed as follows:

*Saccharomyces cerevisiae* and *Saccharomyces paradoxus*:

Yeast Peptone Dextrose (YPD; MP Cat#114001222): 2% Peptone, 1% Yeast extract, 2% Glucose, 2% Agar.

URA dropout: 6.8 g/L Yeast Nitrogen Base (YNB; Sigma-Aldrich Cat#Y0626), 20 g/L Glucose (Sigma-Aldrich Cat#G8270), 0.8 g/L URA dropout (MP Cat#4511-222), 20 g/L Agar, pH 6∼6.5 (with NaOH).

YPD + G418 (Sigma-Aldrich Cat#A1720-1G, 150 mg/L)

YPD + Rapamycin (Selleck Cat#S1039,0.025 μg/mL)

*Schizosaccharomyces pombe*:

Yeast Extract Supplemented (YEA): 30 g/L Glucose, 5 g/L Yeast extract, 100 mg/L Adenine, pH 5.5.

YEA + G418 (Beijing LabLead Biotech Cat#G4180-5G, 100 mg/L)

*Neurospora crassa*:

Minimal slants: 15 g/L Agar, 30 g/L Sucrose, 50**×**Vogel’s.

Bottom Agar/Top Agar: 15 g/L Agar, 10×Fig’s, 50**×**Vogle’s.

Bottom Agar + Hygromycin B (Roche Cat#10843555001, 150 μg/mL).

10**×**Fig’s: 200 g/L L-Sorbose, 5 g/L Glucose, 5 g/L Fructose.

50**×**Vogle’s (1L):

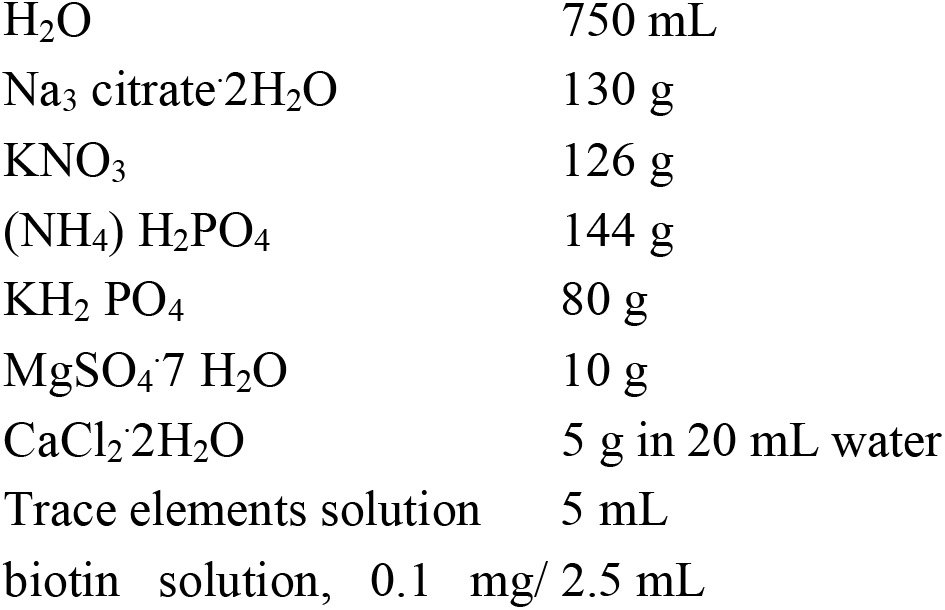
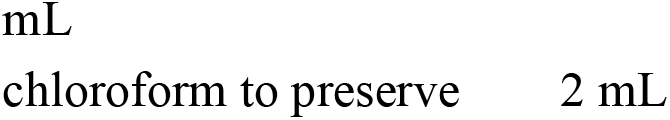

Trace elements solution (100 mL):

|  |  |
| --- | --- |
| H<sub>2</sub>O | 95 mL |
| citric acid.H<sub>2</sub>O | 5 g |
| ZnSO<sub>4</sub>.7H<sub>2</sub>O | 5 g |
| Fe (NH<sub>4</sub>)<sub>2</sub>(SO<sub>4</sub>)<sub>2</sub>.6H<sub>2</sub>O | 1 g |
| CuSO<sub>4</sub>.5H<sub>2</sub>O | 250 mg |
| MnSO<sub>4</sub>.H<sub>2</sub>O | 50 mg |
| H<sub>3</sub>BO<sub>3</sub> | 50 mg |
| Na<sub>2</sub>MoO<sub>4</sub>.2H<sub>2</sub>O | 50 mg |

### Primers

GetPrimers was used to design gene targeting and verification primers for our application demonstrations in *Saccharomyces cerevisiae*, *Saccharomyces paradoxus*, and *Schizosaccharomyces pombe* and partially used to design gene targeting primers for our application in *Neurospora crassa*. The details of these primer combinations are listed as follows.

*Saccharomyces cerevisiae FPR1* gene knockout with the pYES2-*URA3* plasmid in the strains BY4741, DBVPG6044, Y12, YPS128, and UWOPS03-461.4:

P1: ACCTAAACTCGAGTATAAGCAAAAAATCAATCAAAACAAGTAATAAGCTTTTCAATTCAATTCAT

P2: ATAAATAAAAAGCAGAAAGGCGGCTCAATTGATAGTACTTTGCTTTTGAATACTCATACTCTTCC

V1: ATCATCGTCGTTGCTTCTCG

V2: CGAAGAAGGTGATGATGAGCG

V3: CAACCCTTGATGACTTGGCC

V4: GGCCAAGTCATCAAGGGTTG

V5: ATGAATTGAATTGAAAAGCT

V6: GGAAGAGTATGAGTATTCAA

*Saccharomyces paradoxus FPR1* gene knockout with the pBS10-*hphMX* plasmid in the strain CBS432:

P1: ACCTAAACTCGAATATAAGCAAGTAATCAATCAAAAGAAGTAATACGCCAGATCTGTTTAGCTTG

P2: TAAATAAAAAGCAAAAGGCAGCTCAATTGGGGTGGTGTGTTAATTGCTCGTTTTCGACACTGGAT

V1: CCTTACTTGGTGGACTGTGTC

V2: ACGACGGCTCCAGTAAGAAA

V3: CCAACCCTTGATGACTTGGC

V4: GGCCAAGTCATCAAGGGTTG

V5: CAAGCTAAACAGATCTGGCG

V6: ATCCAGTGTCGAAAACGAGC

*Saccharomyces paradoxus FPR1* gene knockout with the pBS10-*hphMX* plasmid in the strain YPS138:

P1: CACCTAAACTCAAATATAAGCAAGTAATAATCAAAAGAAGTAATACGCCAGATCTGTTTAGCTTG

P2: TAAATAAAAAGCAAAAAGCAGCTCAATGGGGGTAGTATCTTAATTGCTCGTTTTCGACACTGGAT

V1: GCTATTGAAAGAGTACGGTGCC

V2: ACAAGTAAAACGACGGCTCC

V3: AATGGAGAGCCCCTGTCAAC

V4: GTCGGACAAGTCATCAAGGG

V5: CAAGCTAAACAGATCTGGCG

V6: ATCCAGTGTCGAAAACGAGC

*Saccharomyces cerevisiae COX4* gene C’-GFP-tagging with the pYES2-*hphMX* plasmid in the strain BY4741:

P1: TACAAACTAAACCCTGTTGGTGTTCCAAATGATGACCACCATCACCGGATCCCCGGGTTAATTAA

P2: GGTAAAAAGTAAAAGAGAAACAGAAGGGCAACTTGAATGATAAGAGAATTCGAGCTCGTTTAAAC

V1: AACAGATCGAACGAAACGGA

V2: GAGAACGAAGCGGTGTAAGC

V5: TTAATTAACCCGGGGATCCG

V6: GTTTAAACGAGCTCGAATTC

*Schizosaccharomyces pombe Ras1* gene knockout with the pFA6a-*kanMX* plasmid in the strain SPJ649:

P1: AGTGTGCTTTATAAAATAGTGAATCGGATCCCCGGGTTAATTAA

P2: CAGCTCAAATATTCTGCAATAATACTTGGAATTCGAGCTCGTTTAAAC

G1: GCCGACCTCTTCATGCATG

G2: AGAAGCCGGTATGAGAGGTT

G3: TTAATTAACCCGGGGATCCGATTCACTATTTTATAAAGCACACT

G4: GTTTAAACGAGCTCGAATTCCAAGTATTATTGCAGAATATTTGAGCTG

V1: TCTTGAGAAACTACATCCTTAACGT

V2: TTTGCAGGAGTTTGTGGTCA

V3: GCGCACCTTCACCATCAATT

V4: GGTGAAGGTGCGCTTTTAGA

V5: TTAATTAACCCGGGGATCCG

V6: GTTTAAACGAGCTCGAATTC

*Schizosaccharomyces pombe Ras1* C’-flag with the pFA6a-3flag-*kanMX* plasmid in the strain SPJ649:

P1: GTTTCAACTAAATGTTGTGTTATATGTCGGATCCCCGGGTTAATTAA

P2: CAGCTCAAATATTCTGCAATAATACTTGGAATTCGAGCTCGTTTAAAC

G1: AATTGATGGTGAAGGTGCGC

G2: AGAAGCCGGTATGAGAGGTT

G3: TTAATTAACCCGGGGATCCGACATATAACACAACATTTAGTTGAAAC

G4: GTTTAAACGAGCTCGAATTCCAAGTATTATTGCAGAATATTTGAGCTG

V1: CAAATTGGTAGTTGTAGGAGATGG

V2: TGCTGGTATGTCGTTTCTTGC

V5: TTAATTAACCCGGGGATCCG

V6: GTTTAAACGAGCTCGAATTC

*Neurospora crassa PRP5* gene (*NCU08674*) knockout with the pCSN44-*hphMX* plasmid in the strain *Ku70^RIP^*:

G1 (*Not*I): AAA<u>GCGGCCGC</u>ACGCATCCTACCTTTCTCCC

G2 (*Kpn*I): CGG<u>GGTACC</u>GTTCATCCATCAACCGAGCC

G3 (*Hind*III): CCC<u>AAGCTT</u>TAGGAGACACCTGGGGAGGAGG

G4 (*Sph*I): ACAT<u>GCATGC</u>GCAGGTAGTCCCTCCCG

hph1: GAAAAAGCCTGAACTCACCG

hph2: CGTCGGTTTCCACTATCGGC

The primer oligonucleotides for the gene targeting experiments in *Saccharomyces cerevisiae* and *Saccharomyces paradoxus* were synthesized by Sangon Biotech (Shanghai) Co., Ltd (Shanghai, China). The primer oligonucleotides for the gene targeting experiments in *Schizosaccharomyces pombe* were synthesized by Beijing Tsingke Biotech Co., Ltd. (Beijing, China). The primers used in the gene targeting experiments for *Neurospora crassa* were synthesized by Beijing Tsingke Biotech Co., Ltd. (Beijing, China).

### Preparation of the DNA fragment for transformation

#### *Saccharomyces cerevisiae* and *Saccharomyces paradoxus*

To construct the *FPR1* knockout strains, the DNA fragment for transformation were created by amplifying the immediate 45-bp upstream and downstream sequences of the targeted gene (*FPR1*) together with the selection marker from the pYES2-*URA3* (*Saccharomyces cerevisiae*) or the pBS10-*hphMX* (*Saccharomyces paradoxus*) plasmid using the P1-P2 primer pair. Such PCR amplifications (25-μL reaction) were performed with the following program: 95°C 5 min for initial denaturation; 95°C 30 sec, 56°C 30 sec, and 72°C 1 min 10 sec (*Saccharomyces cerevisiae*) or 1min 20 sec (*Saccharomyces paradoxus*) for 35 cycles; 72°C 5min for the final extension. To construct the *COX4*-GFP C’-tagged strain, the DNA fragment for transformation was prepared in a similar way. The major alterations are that the upstream flanking sequence was located right upstream of the stop codon of the targeted gene (*COX4*), and the pBS10-*GFP*-*hphMX* plasmid was utilized as the template to amplify the selection marker. Such PCR amplifications (25-μL reaction) were performed with the following program: 95°C 5 min for initial denaturation; 95°C 30 sec, 56°C 30 sec, and 72°C 1 min 10 sec for 35 cycles; 72°C 5min for the final extension. The Premix Taq™ (Takara Cat#R004A) was used for these PCR experiments.

#### Schizosaccharomyces pombe

To construct the *Ras1* knockout strains, DNA fragments for transformation were first created by amplifying the ∼400-bp flanking sequences of the targeted gene (*Ras1*) using the G1-G3 (upstream of the start codon) and G4-G2 (downstream of the stop codon) primer pairs. The *kanMX* selection marker was subsequently amplified from the pFA6a-*kanMX* plasmid using the P1-P2 primer pair. These PCR amplifications (50-μL reaction **×** 4) were performed with the following program: 95°C 3 min for initial denaturation; 95°C 15 sec, 54°C 15 sec, and 72°C 1min (G1-G3 and G4-G2) or 2 min (P1-P2) for 30 cycles; 72°C 5 min for the final extension. A secondary fusion PCR reaction (50-μL reaction **×** 4) was then conducted to combine the flanking fragments and the selection marker fragment generated above (template ratio: 1:1:2) with the G1-G2 primer pair for the deliver of the final DNA fragment for transformation. This fusion PCR was performed with the following program: 95°C 3 min for initial denaturation; 95°C 15 sec, 54°C 15 sec, and 72°C 2 min for 30 cycles; 72°C 5 min for the final extension. The Phanta Max Super-Fidelity DNA Polymerase (Vazyme Cat#P505) was used for these PCR experiments. The preparation of the DNA fragment for the *Ras1*-FLAG strains was performed in a similar way. The major alterations are that the upstream flanking sequence was located right upstream of the stop codon, and the pFA6a-3flag-*kanMX* plasmid was utilized as the template to amplify the selection marker.

#### Neurospora crassa

To generate the *PRP5* gene knockout strain in *Neurospora crassa*, approximately 500-bp flanking sequences of the targeted *PRP5* gene were amplified from genomic DNA extracted from the wild-type strain. The primer pairs G1-G3 (upstream of the start codon) and G4-G2 (downstream of the stop codon) were used to amplify the upstream and downstream fragments, respectively. PCR amplification was performed in a 50-μL reaction with the following program: 98°C 4 min for initial denaturation; 98°C 30 sec, 58°C 30 sec, and 72°C 30 sec for 30 cycles; 72°C 5 min for the final extension. The Gloria Nova HS 2X Master Mix (ABclonal Cat#RK20717) was utilized for these PCR experiments. The upstream fragment was inserted into pCSN44-*hphMX* vector at the *Not*I and *Hind*III sites, resulting in the construct pCSN44-Up-*PRP5*. Subsequently, the downstream fragment was inserted into this vector at the *Sph*I and *Kpn*I sites to generate pCSN44-Up-*Prp5*-Down construct.

### Transformation

#### *Saccharomyces cerevisiae* and *Saccharomyces paradoxus*

The budding yeast transformation was performed following the previously described method [Gietz and Schiestl 2007]. Briefly, the wildtype *Saccharomyces cerevisiae* or *Saccharomyces paradoxus* strain was precultured in flask with YPD medium at 30°C for overnight and then diluted for 25 folds in fresh YPD and cultured for another 5 hours. Cells were harvested after centrifugation and washed twice with sterile water. Cells were then resuspended and washed with 0.1 M LiAc before treated with a master mix containing 50% (w/v) PEG, LiAc (1 M), the PCR product for transformation, and Salmon sperm DNA (2.5 mg/ml). The final mix was vortexed for 2 min before incubation at 30°C for 30 min and 42°C for another 30 min. Cells were collected after centrifugation (8000 rpm **×** 2 min) and washed with sterile water. For cells transformed using auxotrophic markers (e.g., *URA3*), we plated them directly on selection medium plates (e.g., URA dropout). For cells transformed using antibiotic markers (e.g., *kanMX*), we incubate them overnight in YPD before plating onto plates with proper antibiotics (e.g., YPD+G418). The replated cells were incubated at 30°C for 3-4 days before transformant selection.

#### Schizosaccharomyces pombe

The fission yeast transformation was performed following previously described method [Bähler et al. 1998]. Briefly, the wildtype *Schizosaccharomyces pombe* strain was precultured in flask with the YEA medium at 30°C for overnight and then diluted until its OD595 value reached 0.25-0.5. Then the yeast cells were centrifuged and harvest at 4 °C. Cells were resuspended and washed in ice-cold sorbitol (1.2 M) twice. The cuvette was pre-cooled on ice before transformation. Cells were resuspended with sorbitol and further mixed with the PCR product (∼2 μg DNA/200 μL aliquots) for transformation. Immediate pulse at 2.0 kV with MicroPulser Electroporator (BioRad, California, USA) was used for transformation. After transformation, cells were immediately but gently resuspended with ice-cold sorbitol. Cells were first plated on YEA plate for overnight and then replicated onto plates with proper antibiotics (e.g., YEA+G418). The replated cells were incubated at 30°C for 3-4 days before transformant selection.

#### Neurospora crassa

The *Neurospora* transformation was performed as described previously [Zhou, Liu, Hu, Zhang, Sun, Cha, Wang, Liu, and He 2013]. Briefly, the pCSN44-Up-*PRP5*-Down construct was introduced into the host *Ku70^RIP^*strain through electroporation. The positive transformants, with the *hph* gene at the *PRP5* locus, were initially selected by inoculating onto slants containing hygromycin and were subsequently verified by using PCR analyses.

### Spotting assay

Cells were precultured in YPD overnight. Upon saturation, an aliquot of 3.5 μL culture was taken for spotting assay in the condition of interest, for which five 1:10 dilutions were implemented on the plate (from left to right).

### Fluorescence Imaging

For microscopy, 1 mL of yeast cells at exponential phase were collected by centrifugation, washed in PBS and re-suspended in 4% polyformaldehyde (Biosharp Cat#BL539A) to preserve mitochondrial morphology. After 20 min, cells were washed in PBS, and were applied to poly-D-lysine (Sigma-Aldrich Cat# P1149) coated microscope culture dish for 20 min at room temperature. After one more washing step with PBS, cells were visualized using spinning disk confocal (Nikon CSU-W1) equipped with a Plan Apo λ 100**×** oil objective and 488 nm laser lines.

### Western blot

Cells were lysed with 100 μL of cold lysis buffer (50 mM HEPES-KOH, 140 mM NaCl, 1% Triton X-100, 1 mM PMSF) and 1.9 g cold glass beads. The tubes were bead-beaten at top speed for four times (2 min each) and were then inverted to make beads drop to the bottom. The tubes, which were punched in the bottom,were placed into a 5 mL collection tube. The tube was centrifuged at 1500 rpm for 2 min at 4°C. After vortex, the lysates were carefully transferred to an eppendorf tube and centrifuged at 13,000 rpm for 15 min at 4°C.The extracted protein was loaded to SDS-PAGE gel for electrophoresis at 110 V for 1 hour. Proteins were then transferred to a nitrocellulose membrane using the Bio-Rad Trans-Blot Turbo transfer system. Membranes were blocked for 1 hour at room temperature in blocking buffer (Beyotime Cat#P0252-500-mL). Then membranes were incubated with 1:5000 diluted primary antibodies (Sigma-Aldrich Cat#F3165) in TBST-5% nonfat dried milk overnight at 4°C. Membranes were washed three times (10 min each) in TBST. The membranes were incubated with HRP-conjugated secondary antibody at 1:10,000 in TBST-5% nonfat dried milk for 1 hour at room temperature. For chemiluminescence, membranes were washed 3 times (10 min each). Protein signals were visualized with Azure Biosystems C600 image analyzer (Dublin, California, USA)

## Results and Discussion

### The algorithm design of GetPrimers

GetPrimers is a generalized computational tool developed for automatic and fully customizable primer design of PCR-based gene targeting. While it can also be deployed and used locally via command lines, we provide a dedicated web server (https://www.evomicslab.org/app/getprimers/) to facilitate researchers to conveniently design PCR-based gene targeting and verification primers by only a few clicks. GetPrimers natively supports gene targeting based on both one-step PCR (the “long primer” strategy) and fusion PCR (the “short primer” strategy), making it highly amenable for diverse use scenarios.

For the one-step-PCR-based strategy, a single pair of long PCR primers named P1 and P2 are automatically determined based on the selection marker cassette of the targeting plasmid as well as the 5’- and 3’-flanking regions of the targeted genomic locus. The corresponding PCR fragment will be swapped into the targeted locus guided by the locus-specific sequences designed on P1 and P2 (Figure 1A). For fusion-PCR-based strategy, three pairs of short PCR primers (namely P1-P2, G1-G3, and G2-G4) are automatically picked for amplifying fragments corresponding to the selection marker cassette, the 5’-flanking homologous arm and the 3’-flanking homologous arm respectively, which will be further fused into a larger fragment for target incorporation guided by the amplified homologous arms (Figure 1B). For both one-step-PCR-based and fusion-PCR-based gene targeting, a suite of verification primers will be further designed (Figure 2) to help researchers to check and pick transformed clones with correct on-target integrations. The designed gene targeting primers and verification primers are automatically scored, ranked, and filtered based on various considerations such as genome-wide specificity, dimer formation, etc.

**Figure 1.**
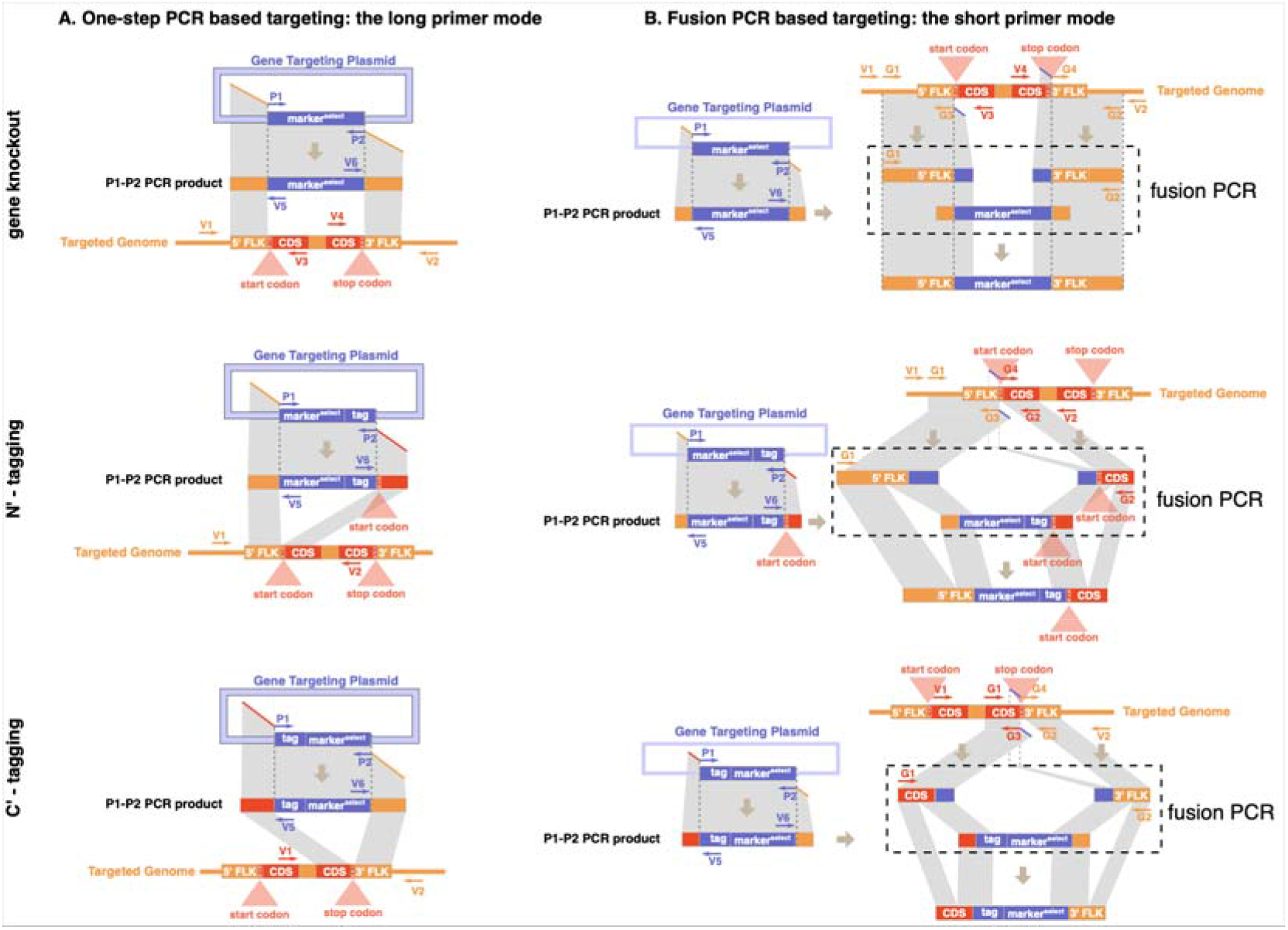
The targeting and verification primer design of GetPrimers. (A) The primer design for one-step-PCR-based targeting strategy. (B) The primer design for fusion-PCR-based targeting strategy. Primers denoted as P1, P2, and G1-G4 are gene targeting primers. Primer denoted as V1-V6 are verification primers. CDS: coding sequence. 5’ FLK: 5’-flanking sequence. 3’ FLK: 3’ flanking sequence.

**Figure 2.**
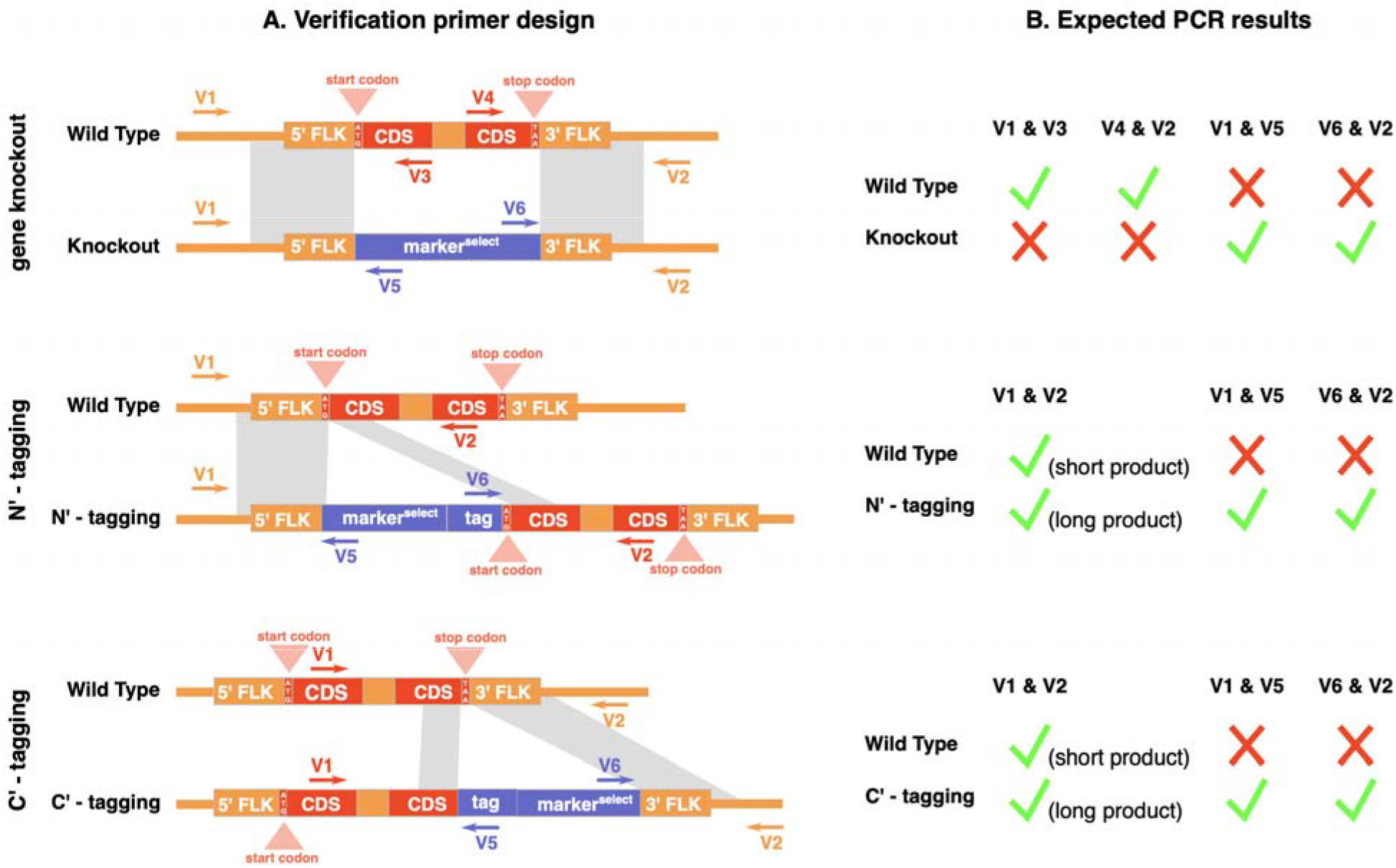
The design and expected PCR results of verification primers. (A) The genomic location of verification primers both before (wild type) and after (knockout, N’-tagging, and C’-tagging) the corresponding gene targeting experiment. (B) The expected PCR result of each verification primer combination for both untargeted and targeted clones. The green check mark denotes for expecting to have PCR product. The red cross mark denotes for expecting to have no PCR product.

GetPrimers was implemented in Perl and bash at the backend, with its frontend web portal powered by the python-based Django framework. Primer3 was used to facilitate primer searching, assessment, and ranking [Koressaar and Remm 2007]. With GetPrimers’ web portal, we have pre-computed targeting and verification primers for all protein-coding genes of several representative yeast and fungi genomes (e.g., *Saccharomyces cerevisiae*, *Saccharomyces paradoxus, Schizosaccharomyces pombe*, and *Neurospora crassa*). Commonly used gene targeting plasmid backbones such as pFA6a [Bähler et al. 1998] and pYES2 [Kevin R., Vo, Michaelis, and Paddon 1997] were adopted for such genome-wide calculation. The pre-computed results were stored in an SQL3 database. Meanwhile, the web portal also allows user to submit their own input genome and targeting plasmid sequences for fully customized targeting and verification primer design.

### Application demonstrations

To demonstrate the usage and versatility of GetPrimers, we applied it to various gene targeting experiments using the budding yeast *Saccharomyces cerevisiae* and *Saccharomyces paradoxus*, the fission yeast *Schizosaccharomyces pombe*, as well as the filamentous fungus *Neurospora crassa* (Table 3). Successful constructed strains were obtained for all these experiments with relatively high successful rates. These constructed strains were all verified by PCR using the verification primers designed by GetPrimers. Phenotyping-based verification were further performed for those with additional observable phenotypes, such as acquiring Rapamycin resistance when deleting *FRP1*, exhibiting fluorescence when tagged with green fluorescent protein (GFP), and showing up in western blot when tagged with FLAG.

**Table 3.** A summary of application demonstration experiments performed for GetPrimers.

| Species | Strain | Ploidy | Targeting experiment type | Targeted gene | Targeting plasmid backbone | Selection marker | Successful ratio |
| --- | --- | --- | --- | --- | --- | --- | --- |
| *Saccharomyces cerevisiae* | BY4741 | 1n | one-step-PCR-based knockout | *FPR1* | pYES2 | *URA3* | 1/12 |
|  | DBVPG6044 | 1n | one-step-PCR-based knockout | *FPR1* | pYES2 | *URA3* | 2/8 |
|  | Y12 | 1n | one-step-PCR-based knockout | *FPR1* | pYES2 | *URA3* | 3/16 |
|  | YPS128 | 1n | one-step-PCR-based knockout | *FPR1* | pYES2 | *URA3* | 2/16 |
|  | UWOPS03-461.4 | 1n | one-step-PCR-based knockout | *FPR1* | pYES2 | *URA3* | 1/8 |
| *Saccharomyces paradoxus* | CBS432 | 2n | one-step-PCR-based knockout | *FPR1* | pBS10 | *hphMX* | 16/16 |
|  | YPS138 | 2n | one-step-PCR-based knockout | *FPR1* | pBS10 | *hphMX* | 13/16 |
| *Saccharomyces cerevisiae* | BY4741 | 1n | one-step-PCR-based C'-GFP-tagging | *COX4* | pFA6a | *hphMX* | 14/14 |
| *Schizosaccharomyces pombe* | SPJ649 | 1n | fusion-PCR-based knockout | *Ras1* | pFA6a | *kanMX* | 22/24 |
| *Schizosaccharomyces pombe* | SPJ649 | 1n | fusion-PCR-based C'-FLAG-tagging | *Ras1* | pFA6a | *kanMX* | 23/24 |
| *Neurospora crassa* | *Ku70<sup>RIP</sup>* | 1n | fusion-PCR-based knockout | *PRP5* | pCSN44 | *hphMX* | 4/9 |

### Application demonstration 1: one-step-PCR-based knockout of the *FPR1* gene in the budding yeast *Saccharomyces cerevisiae*

The *FPR1* gene in the budding yeast *Saccharomyces cerevisiae* encodes the Peptidyl-prolyl cis-trans isomerase (PPIase), which can bind to Rapamycin, a commonly used anti-tumor chemotherapy drug. Our previous study in *Saccharomyces cerevisiae* showed that the loss of function mutation of this gene can lead to substantial cellular resistance to Rapamycin across multiple genetic backgrounds [Li et al. 2019]. Here we used GetPrimers to design primers for one-step-PCR-based *FPR1* knockout across five *Saccharomyces cerevisiae* strains: BY4741, DBVPG6044, Y12, YPS128, UWOPS03-461.4. Among these five strains, BY4741 is a commonly used lab strain, whereas the other four strains are representative natural strains isolated from diverse geographical locations: West Africa for DBVPG6044, Japan for Y12, North America for YPS128, and Malaysia for UWOPS03-461.4 [Liti et al. 2009; Yue et al. 2017]. Given the relative moderate population polymorphism level (0.5%) (Supplementary Table 1) and high homologous recombination efficiency of *Saccharomyces cerevisiae*, we used the same targeting and verification primer combination designed by GetPrimers based on the *Saccharomyces cerevisiae* S288C reference genome for this experiment. With a selection marker of *URA3*, we managed to construct the *fpr1*Δ strains based on all five haploid strain backgrounds with a successful rate of 8.3% - 25% as suggested by our verification PCRs (Table 3 & Figure 3A-D). Moreover, since the deletion of the *FPR1* gene will induce Rapamycin resistance, we performed the spotting assay on Rapamycin-containing medium (0.025 µg/mL) to further validate these successful constructs at the phenotypic level (Figure 3E).

**Figure 3.**
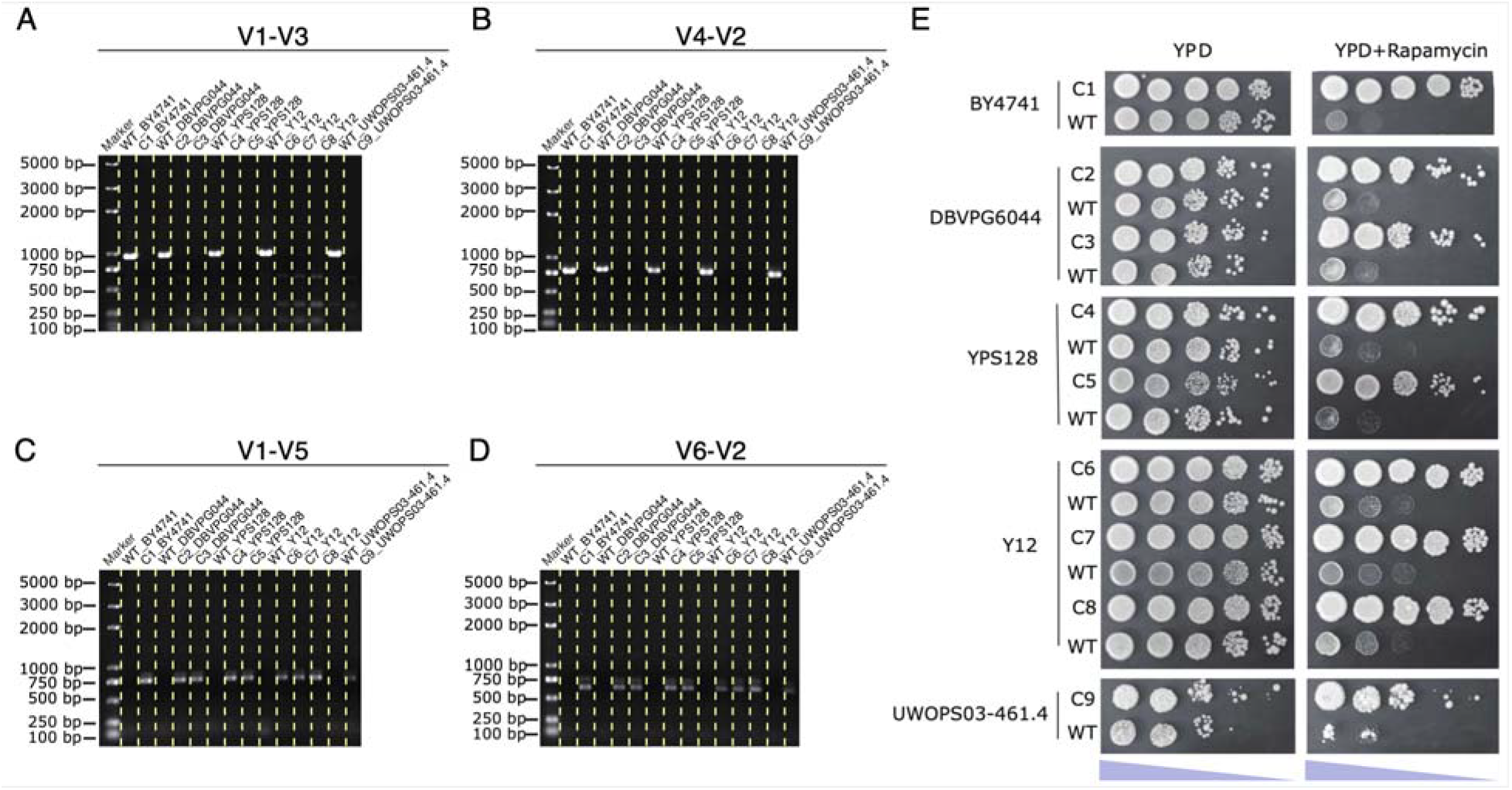
The PCR and phenotyping validation of the *FPR1* deletion in *Saccharomyce cerevisiae*. (A) The PCR validation results using the verification primer V1 & V3. (B) The PCR validation results using the verification primer V4 & V2. (C) The PCR validation results using the verification primer V1 & V5. (D) The PCR validation results using the verification primer V6 & V2. (E) The phenotyping validation results based on Rapamycin (0.025 µg/mL with 10**×** serial dilution) resistance readouts. WT: wild type. C1-C9: validated clones with correct *FPR1* gene deletion.

### Application demonstration 2: one-step-PCR-based knockout of the *FPR1* gene in the budding yeast *Saccharomyces paradoxus* with highly divergent genetic backgrounds

As the closest sister species of *Saccharomyces cerevisiae*, *Saccharomyces paradoxus* has a much more stratified population structure with substantial subpopulation separations [Liti et al. 2009; Yue et al. 2017]. Such higher polymorphism levels between subpopulations means common targeting and verification primers designed based on a single reference will be less efficient for strains from other subpopulations. Instead, background-specific primer design should warrantee a more efficient gene targeting experiment. Therefore, we let GetPrimers to design background-specific primers for deleting the *FPR1* gene with one-step PCR in two *Saccharomyces paradoxus* strains with highly divergent genetic backgrounds: the Far East strain CBS432 and the North American strain YPS138 (Supplementary Figure 1). A highly efficient *FPR1* knockout was achieved for both diploid CBS432 and YPS138 strains as evaluated by our PCR verification, with a successful rate of 100% and 81.2% respectively (Table 3). Given that the CBS432 and YPS138 strains that we used here are diploid strains, normally their *FPR1* knockout will be heterozygous (Figure 4A-B). Interestingly, in one case (C1 from the *FPR1* knockout in the CBS432 background), however, we found the deletion to be homozygous, hinting the occurrence of a secondary loss of heterozygosity (LOH) event. Accordingly, we also observed a stronger Rapamycin resistance for this C1 strain compared with its counterparts with heterozygous *FPR1* deletion in the same CBS432 background (Figure 4C-D).

**Figure 4.**
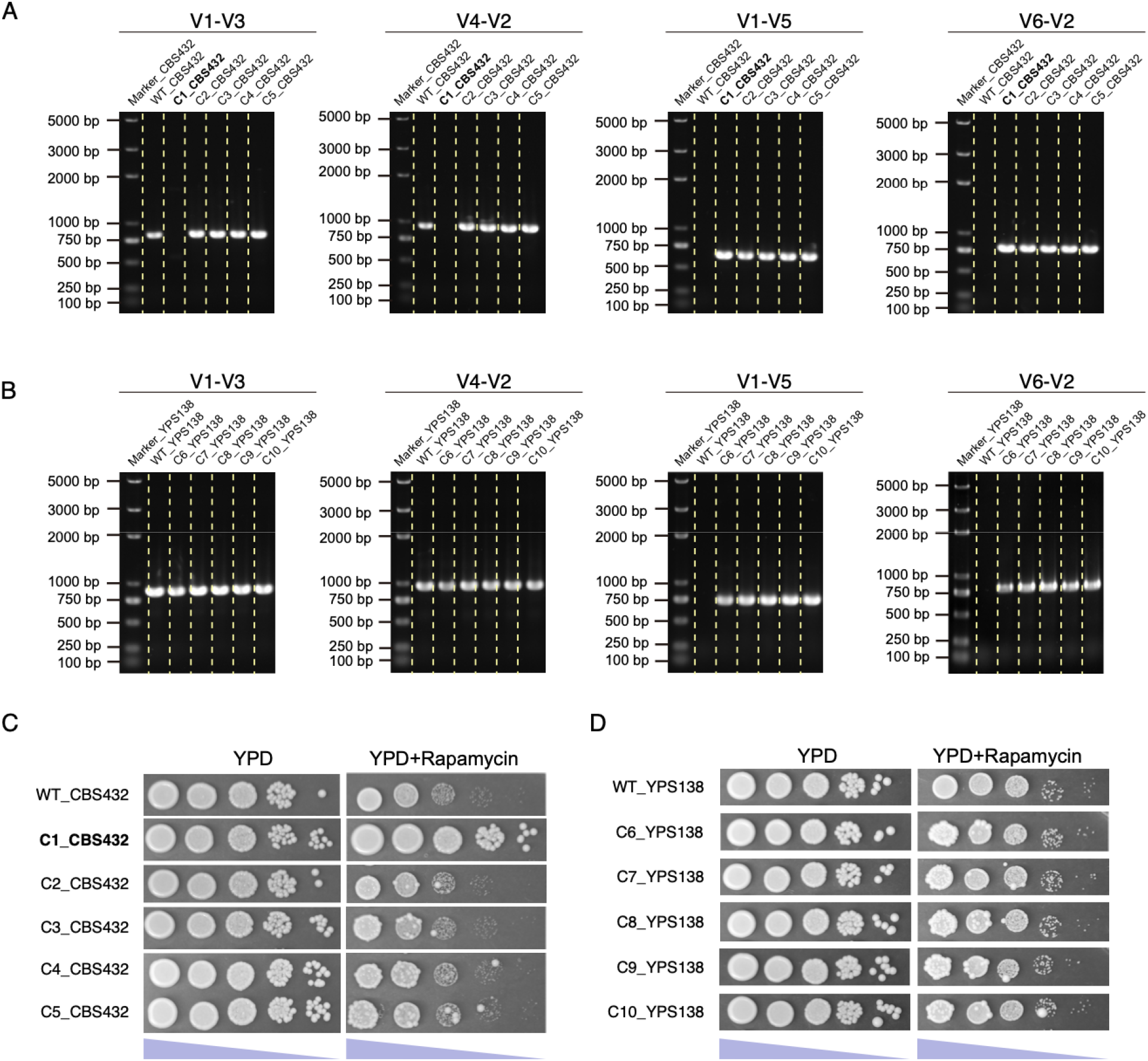
The PCR and phenotyping validation of the *FPR1* deletion in *Saccharomyces paradoxus*. (A-B) The PCR validation results for CBS432-based (A) and YPS138-based (B) *FPR1* deletion using the verification primer V1 & V3, V4 & V2, V1 & V5, and V6 & V2. (C-D) The phenotyping validation results for CBS432-based (C) and YPS138-based (D) *FPR1* deletion based on Rapamycin (0.025 µg/mL with 10**×** serial dilution) resistance readouts. WT: wild type. C1-C10: validated clones with correct *FPR1* gene deletion. The clone with homozygous *FPR1* deletion (C1_CBS432) was denoted in bold.

### Application demonstration 3: the one-step-PCR-based C’-GFP-tagging of the *COX4* gene in the budding yeast *Saccharomyces cerevisiae*

In *Saccharomyces cerevisiae*, the *COX4* gene locates in the nuclear genome but encode the subunit IV of the cytochrome *c* oxidase that functions in mitochondria as a terminal enzyme of the respiratory chain. Here we want to attach a GFP at the 3’-terminus of the *COX4* gene to form a fusion gene capable of emitting spontaneous fluorescent signals. We used GetPrimers to design the C’-tagging and verification primers for this gene in *Saccharomyces cerevisiae* using one-step PCR strategy. As suggested by our PCR verification, our strain construction achieved an impressive 100% (14/14) successful rate (Table 3). Under the microscope, we observed bright green foci in these constructed yeast cells, which suggested that our GFP-tagging for the *COX4* gene worked as expected.

### Application demonstration 4: fusion-PCR-based knockout and C’-FLAG-tagging of the ***Ras1* gene in the fission yeast *Schizosaccharomyces pombe*.**

The *Ras1* gene in the fission yeast *Schizosaccharomyces pombe* encodes the GTPase Ras protein, which is a homolog of mammalian *RAS* proto-oncogenes [Pylayeva-Gupta, Grabocka, and Bar-Sagi 2011]. Using GetPrimers, we designed primers for fusion-PCR-based gene knockout and C’-FLAG-tagging in *Schizosaccharomyces pombe*. According to our PCR verification, the *Ras1* gene has been precisely targeted with a successful rate > 90% for both knockout and C’-FLAG-tagging experiments (Table 3 and Figure 6A-B). For constructed strains with the C’-FLAG tag, we further applied western blot to demonstrate the expression of the Ras1 protein using the attached FLAG tag (Figure 6C-D).

**Figure 5.**
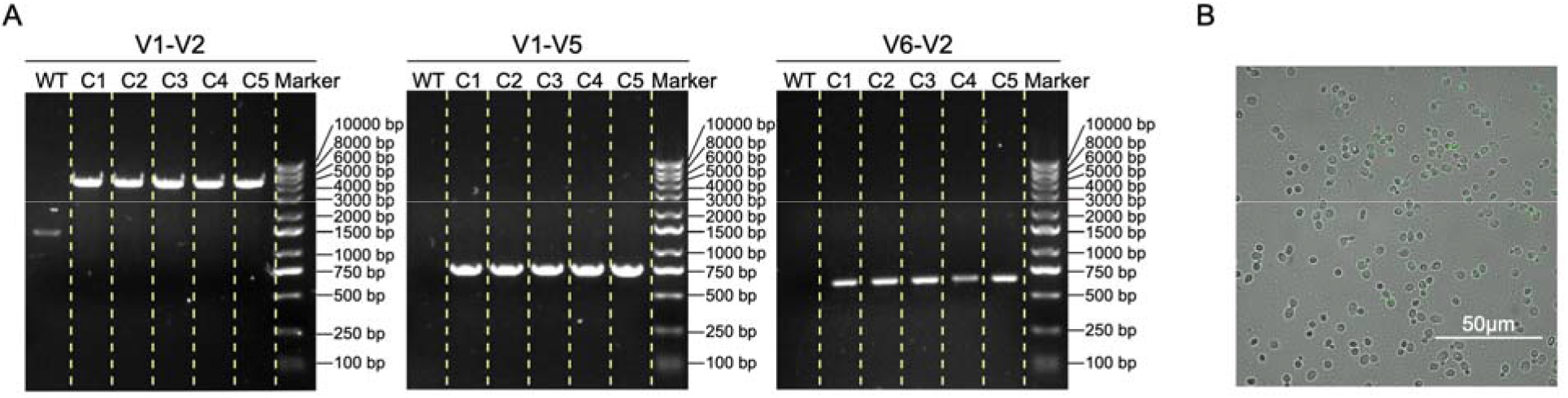
The PCR and imaging validation of the *COX4* C’-GFP-tagging in *Saccharomyce cerevisiae*. (A) The PCR validation results using the verification primer V1 & V2, V1 & V5, and V6 & V2. (B) The fluorescence imaging validation result.

**Figure 6.**
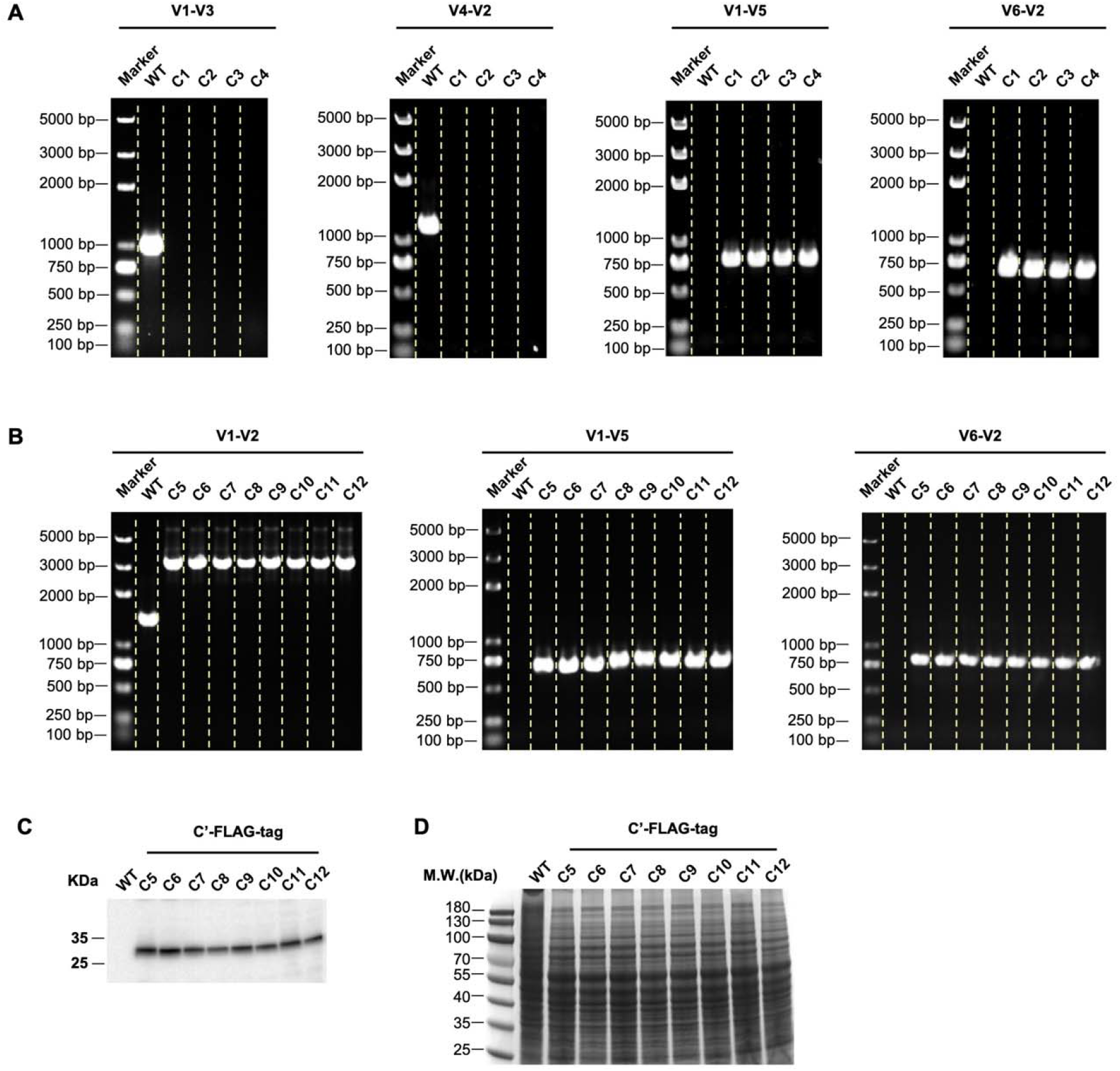
The PCR and western blot validation of the *Ras1* deletion and C’-FLAG tagging in *Schizosaccharomyces pombe*. (A-B) The PCR validation results for *Ras1* deletion using the verification primer V1 & V3, V4 & V2, V1 & V5, and V6 & V2. (B) The PCR validation results for *Ras1* C’-FLAG tagging using the verification primer V1 & V2, V1 & V5, and V6 & V2. (C) The western blot validation results for *Ras1* C’-FLAG tagging. (D) The loaded total protein for the western blot experiment reflected by the Coomassie brilliant blue (CBB) staining. WT: wild type. C1-C4: validated clones with correct *Ras1* gene deletion. C5-C12: validated clones with correct *Ras1* C’-FLAG tagging.

### Application demonstration 5: the adoption of GetPrimers-designed gene targeting primers in vector-clone-based *PRP5* gene knockout in the filamentous fungus *Neurospora crassa*

The filamentous fungus *Neurospora crassa* is an important model eukaryote that has contributed substantially to our understanding of genome defense, circadian clock, meiotic recombination and vegetative incompatibility[Galagan et al. 2003]. Conventionally, gene knockout in *Neurospora crassa* was achieved using a vector cloning-based method. Here we showed how GetPrimers-designed gene targeting primers can be directly applied to vector cloning-based gene knockout. The *PRP5* gene (identified as NCU08674 in *Neurospora crassa*) is a member of the pentatricopeptide repeat protein (PRP) encoding gene family that influences both RNA stability and splicing. First, two PCR products were generated using GetPrimers’ gene targeting primer combinations G1-G3 and G4-G2. These fragments were further integrated into an intermediate vector using enzyme digestion and enzyme ligation. The resulting knockout construct was introduced into the host strain via electroporation and then selected with hygromycin. Our PCR validation revealed that the *PRP5* gene was successfully deleted in 4 out of 9 randomly selected hygromycin-resistant transformants (Figure 7 and Table 3). This result indicates that primers designed by GetPrimers can be utilized to create knockout mutants in *Neurospora crassa* with high efficiency.

**Figure 7.**
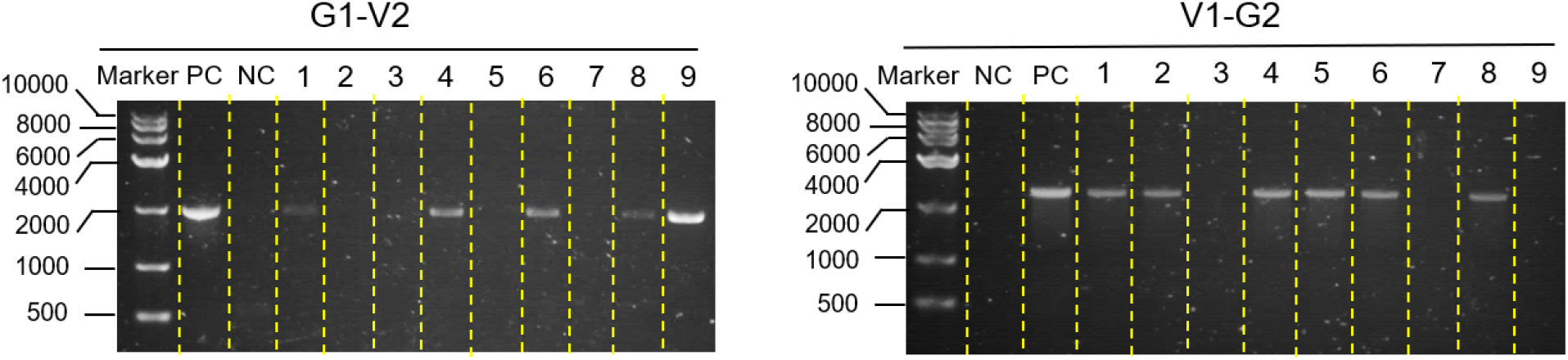
The PCR validation of the *PRP5* gene knockout in *Neurospora crassa*. The PCR verification was conducted using the primer pairs G1 & V2, and V1 & G2. In these analyses, the knockout construct for *PRP5* served as the positive control (PC), while genomic DNA extracted from the wild-type strain was used as the negative control (NC).

## Conclusions

One most powerful technique for studying gene function at the molecular level is to specifically target this gene and perform genetic manipulations such as gene disruption, modification, or tagging. In fungi and many other eukaryotic microorganisms, such gene targeting based on homologous recombination of PCR-products is widely used, which serves to directly manipulate the targeted gene *in situ* without involving any cloning step. This technique is one of a kind in terms of easy gene manipulating before the discovery of CRISPR-Cas9 genome editing system.

In this work, we present GetPrimers, a generalized gene targeting primer designer based on PCR and homologous recombination. In addition to the command line tool implementation, a dedicated webserver with intuitive graphical interface is further provided to make it more accessible for bench researchers. The versatility and adaptability of GetPrimers, supporting both easy-to-use one-step and fusion PCR strategies, makes it a highly valuable resource for researchers working on genetic manipulation in a variety of eukaryotic species. GetPrimers shines in its versatility as it can be applied to any eukaryotic system, any genetic background, and any genomic locus by design. Batch processing is further supported with genome-wide scalability. Therefore, one of the key strengths of GetPrimers is its ability to facilitate large-scale experimental design with full scalability. This feature is especially beneficial for projects that require genome-wide or a large number of gene targeting experiments. Furthermore, GetPrimers can design targeting primers for any user-defined genomic locus, which expands its potential application beyond just protein-coding genes.

As proofs of concept, we performed a serial of gene targeting experiments with primers designed by GetPrimers across multiple systems, including *Saccharomyces cerevisiae*, *Saccharomyces paradoxus*, *Schizosaccharomyces pombe*, and *Neurospora crassa*. The successful application of GetPrimers in various gene targeting experiments, as demonstrated in this study, underscores the tool’s effectiveness and reliability. The high success rates achieved in the construction of strains across multiple yeast and fungi species also attest to GetPrimers’ versatility and robustness. Altogether, we anticipate GetPrimers become a highly useful tool to facilitate easy and standardized genetic targeting and manipulation at scale across multiple eukaryotic systems.

## Acknowledgements

We thank Dr. Li-Lin Du and Dr. Gaowen Liu for helping us to set up the experimental and technical foundation for working with the fission yeasts. We thank the two anonymous reviewers for their highly constructive comments, which help to improve the quality and clarity of our manuscript and the associated software.

## Data availability statement

The command line version of GetPrimers is written in Perl and bash, hosted at https://github.com/codeatcg/getprimers. The web version of GetPrimers is powered by the python-based Django framework, hosted at https://www.evomicslab.org/app/getprimers/. Both versions are free for use under the MIT license. The key raw data are uploaded to the Research Deposit public platform (www.researchdata.org.cn), with the approval RDD number of RDDB2023268940.

## Author contribution statement

**Zepu Miao**: methodology; investigation; data curation; formal analysis; writing—original draft; writing—review and editing. **Haiting Wang**: validation; investigation; data curation; formal analysis; writing—original draft; writing—review and editing. **Xinyu Tu**: validation; investigation; data curation; formal analysis; writing—original draft; writing—review and editing. **Zhengshen Huang**: validation; investigation; data curation; formal analysis; writing— original draft; writing—review and editing. **Shujing Huang**: validation; investigation; data curation; formal analysis; writing—original draft; writing—review and editing. **Xinxin Zhang**: validation; investigation; data curation; formal analysis; writing—original draft; writing—review and editing. **Fan Wang**: validation; investigation; data curation. **Zhishen Huang**: validation; investigation; data curation. **Huihui Li**: investigation; data curation. **Yue Jiao**: resources; investigation; data curation. **Song Gao**: funding acquisition; resources; investigation; data curation. **Zhipeng Zhou**: funding acquisition; resources; methodology; investigation; data curation; writing—original draft; writing—review and editing. **Chun-min Shan**: Conceptualization; supervision; funding acquisition; investigation; methodology; resources; writing—original draft; writing—review and editing. **Jing Li**: Conceptualization; supervision; funding acquisition; investigation; methodology; resources; writing—original draft; writing— review and editing. **Jia-Xing Yue**: Conceptualization; supervision; project administration; funding acquisition; methodology; resources; investigation; data curation; writing—original draft; writing—review and editing.

## References

Amberg DC, Botstein D, Beasley EM. 1995. Precise gene disruption in Saccharomyces cerevisiae by double fusion polymerase chain reaction. Yeast, 11: 1275–1280.

Bähler J, Wu J-Q, Longtine MS, Shah NG, Mckenzie III A, Steever AB, Wach A, Philippsen P, Pringle JR. 1998. Heterologous modules for efficient and versatile PCR-based gene targeting in Schizosaccharomyces pombe. Yeast, 14: 943–951.

Baudin A, Ozier-Kalogeropoulos O, Denouel A, Lacroute F, Cullin C. 1993. A simple and efficient method for direct gene deletion in Saccharomyces cerevisiae. Nucleic Acids Res, 21: 3329–3330.

Camacho C, Coulouris G, Avagyan V, Ma N, Papadopoulos J, Bealer K, Madden TL. 2009. BLAST+: architecture and applications. BMC Bioinformatics, 10: 421.

Colot HV, Park G, Turner GE, Ringelberg C, Crew CM, Litvinkova L, Weiss RL, Borkovich KA, Dunlap JC. 2006. A high-throughput gene knockout procedure for Neurospora reveals functions for multiple transcription factors. Proceedings of the National Academy of Sciences, 103: 10352–10357.

Cummings MT, Joh RI, Motamedi M. 2015. PRIMED: PRIMEr Database for Deleting and Tagging All Fission and Budding Yeast Genes Developed Using the Open-Source Genome Retrieval Script (GRS). PLOS ONE, 10: e0116657.

Cuomo CA et al. 2007. The Fusarium graminearum Genome Reveals a Link Between Localized Polymorphism and Pathogen Specialization. Science, 317: 1400–1402.

Dean RA et al. 2005. The genome sequence of the rice blast fungus Magnaporthe grisea. Nature, 434: 980–986.

Fu J, Hettler E, Wickes BL. 2006. Split marker transformation increases homologous integration frequency in Cryptococcus neoformans. Fungal Genet Biol, 43: 200–212.

Galagan JE et al. 2003. The genome sequence of the filamentous fungus Neurospora crassa. Nature, 422: 859–868.

Giaever G et al. 2002. Functional profiling of the Saccharomyces cerevisiae genome. Nature, 418: 387–391.

Gietz RD, Schiestl RH. 2007. High-efficiency yeast transformation using the LiAc/SS carrier DNA/PEG method. Nat Protoc, 2: 31–34.

Goffeau A et al. 1996. Life with 6000 Genes. Science, 274: 546–567.

Goins CL, Gerik KJ, Lodge JK. 2006. Improvements to gene deletion in the fungal pathogen Cryptococcus neoformans: Absence of Ku proteins increases homologous recombination, and co-transformation of independent DNA molecules allows rapid complementation of deletion phenotypes. Fungal Genetics and Biology, 43: 531–544.

Goldstein AL, McCusker JH. 1999. Three new dominant drug resistance cassettes for gene disruption in Saccharomyces cerevisiae. Yeast, 15: 1541–1553.

Jia G-S, Zhang W-C, Liang Y, Liu X-H, Rhind N, Pidoux A, Brysch-Herzberg M, Du L-L. 2023. A high-quality reference genome for the fission yeast Schizosaccharomyces osmophilus. G3 Genes|Genomes|Genetics, 13: jkad028.

Kevin R. O, Vo KT, Michaelis S, Paddon C. 1997. Recombination-mediated PCR-directed plasmid construction in vivo in yeast. Nucleic Acids Research, 25: 451–452.

Klosterman SJ et al. 2011. Comparative Genomics Yields Insights into Niche Adaptation of Plant Vascular Wilt Pathogens. PLOS Pathogens, 7: e1002137.

Kooistra R, Hooykaas PJJ, Steensma HY. 2004. Efficient gene targeting in Kluyveromyces lactis. Yeast, 21: 781–792.

Koressaar T, Remm M. 2007. Enhancements and modifications of primer design program Primer3. Bioinformatics, 23: 1289–1291.

Kuwayama H, Obara S, Morio T, Katoh M, Urushihara H, Tanaka Y. 2002. PCR-mediated generation of a gene disruption construct without the use of DNA ligase and plasmid vectors. Nucleic Acids Research, 30: e2.

Li J et al. 2019. Shared molecular targets confer resistance over short and long evolutionary timescales. Mol Biol Evol, 36: 691–708.

Liti G et al. 2009. Population genomics of domestic and wild yeasts. Nature, 458: 337–341.

Morgulis A, Gertz EM, Schäffer AA, Agarwala R. 2006. WindowMasker: window-based masker for sequenced genomes. Bioinformatics, 22: 134–141.

Nikawa J, Kawabata M. 1998. PCR- and ligation-mediated synthesis of marker cassettes with long flanking homology regions for gene disruption in Saccharomyces cerevisiae. Nucleic Acids Research, 26: 860–861.

Penkett CJ, Birtle ZE, Bähler J. 2006. Simplified primer design for PCR-based gene targeting and microarray primer database: two web tools for fission yeast. Yeast, 23: 921–928.

Pylayeva-Gupta Y, Grabocka E, Bar-Sagi D. 2011. RAS oncogenes: weaving a tumorigenic web. Nat Rev Cancer, 11: 761–774.

Rhind N et al. 2011. Comparative Functional Genomics of the Fission Yeasts. Science, 332: 930– 936.

Rothstein RJ. 1983. One-step gene disruption in yeast. Methods Enzymol, 101: 202–211.

Taniguti LM et al. 2015. Complete Genome Sequence of Sporisorium scitamineum and Biotrophic Interaction Transcriptome with Sugarcane. PLOS ONE, 10: e0129318.

Untergasser A, Cutcutache I, Koressaar T, Ye J, Faircloth BC, Remm M, Rozen SG. 2012. Primer3—new capabilities and interfaces. Nucleic Acids Research, 40: e115.

Wach A, Brachat A, Pöhlmann R, Philippsen P. 1994. New heterologous modules for classical or PCR-based gene disruptions in Saccharomyces cerevisiae. Yeast, 10: 1793–1808.

Wang X, Xu R, Wang Y, Liu Z, Lou R, Sugiyama T. 2021. Yesprit and Yeaseq: Applications for designing primers and browsing sequences for research using the four Schizosaccharomyces species. Yeast, 38: 583–591.

Wendland J, Ayad-Durieux Y, Knechtle P, Rebischung C, Philippsen P. 2000. PCR-based gene targeting in the filamentous fungus Ashbya gossypii. Gene, 242: 381–391.

Winzeler EA et al. 1999. Functional characterization of the S. cerevisiae genome by gene deletion and parallel analysis. Science, 285: 901–906.

Wood V et al. 2002. The genome sequence of Schizosaccharomyces pombe. Nature, 415: 871– 880.

Yofe I, Schuldiner M. 2014. Primers-4-Yeast: a comprehensive web tool for planning primers for Saccharomyces cerevisiae. Yeast, 31: 77–80.

You B-J, Lee M-H, Chung K-R. 2009. Gene-specific disruption in the filamentous fungus Cercospora nicotianae using a split-marker approach. Arch Microbiol, 191: 615–622.

Yue J-X et al. 2017. Contrasting evolutionary genome dynamics between domesticated and wild yeasts. Nature Genetics, 49: 913–924.

Zhou Z, Liu X, Hu Q, Zhang N, Sun G, Cha J, Wang Y, Liu Y, He Q. 2013. Suppression of WC-independent frequency transcription by RCO-1 is essential for Neurospora circadian clock. Proceedings of the National Academy of Sciences, 110: E4867–E4874.

