## Supplementary Figure 1 for "GetPrimers: a generalized PCR-based genetic targeting primer designer enabling easy and standardized targeted gene modification across multiple systems"

**Supplementary Table 1.** Knockout primers designed by GetPrimers by one-step-PCR strategy for *FPR1* in *Saccharomyces cerevisiae*

| Strains | P1 | P2 |
| --- | --- | --- |
| S288C | ACCTAAACTCGAGTATAAGCAAAAAATCAATCAAAACAAGTAATACGGATCCCCGGGTTAATTAA | ATAAATAAAAAGCAGAAAGGCGGCTCAATTGATAGTACTTTGCTTGAATTCGAGCTCGTTTAAAC |
| BY4741 | ACCTAAACTCGAGTATAAGCAAAAAATCAATCAAAACAAGTAATACGGATCCCCGGGTTAATTAA | ATAAATAAAAAGCAGAAAGGCGGCTCAATTGATAGTACTTTGCTTGAATTCGAGCTCGTTTAAAC |
| DBVPG6044 | ACCTAAACTCGAATATAAGCAAAAAATCAATCAAAACAAGTAATACGGATCCCCGGGTTAATTAA | ATAAATAAAAAGCAGAAAGGCGGCTCAATTGATAGTACTTTGCTTGAATTCGAGCTCGTTTAAAC |
| Y12 | ACCTAAACTCGAATATAAGCAAAAAATCAATCAAAACAAGTAATACGGATCCCCGGGTTAATTAA | ATAAATAAAAAGCAGAAAGGCGGCTCAATTGATAGTACTTTGCTTGAATTCGAGCTCGTTTAAAC |
| YPS128 | ACCTAAACTCGAATATAAGCAAAAAATCAATCAAAACAAGTAATACGGATCCCCGGGTTAATTAA | ATAAATAAAAAGCAGAAAGGCGGCTTAATTGATAGTACTTTGCTTGAATTCGAGCTCGTTTAAAC |
| UWOPS03-461.4 | ACCTAAACTCGAATATAAGCAAAAAATCAATCAAAACAAGTAATACGGATCCCCGGGTTAATTAA | ATAAATAAAAAGCAGAAAGGCGGCTCAATTGATAGTACTTTGCTTGAATTCGAGCTCGTTTAAAC |

Note: the polymorphic bases compared with the *Saccharomyces cerevisiae* S288C reference are highlighted.

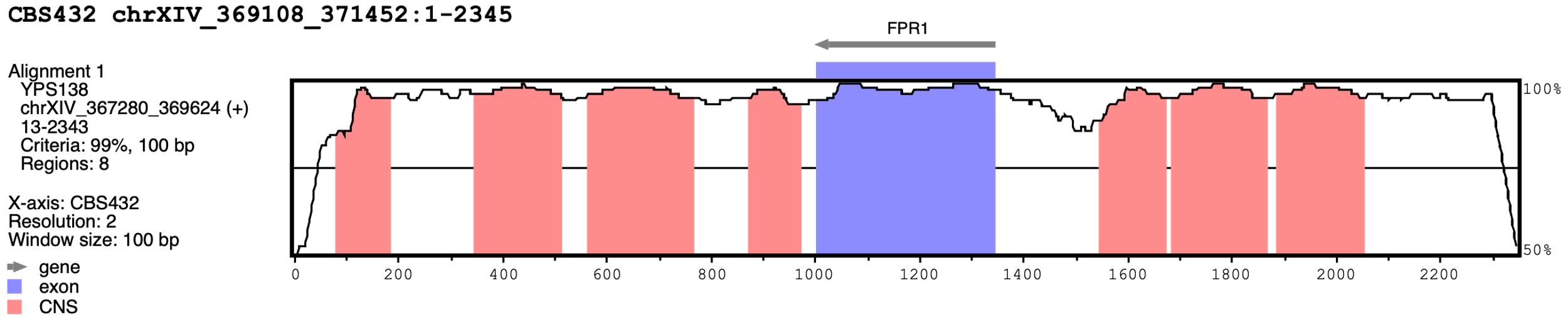

**Supplementary Figure 1.** The sequence conservation profile of *FPR1* and its flanking regions between *Saccharomyces* *paradoxus* CBS432 and *Saccharomyces* *paradoxus* YPS138. The sequence identity between CBS432 and YPS138 is 99.1% for the *FPR1* coding region (1001 – 1345 bp) while being 95.3% (1 – 1000 bp) and 96.0% (1346 – 2345 bp) for the *FPR1*’s upstream and downstream flanking regions respectively. The plot was generated by mVISTA (https://genome.lbl.gov/vista/mvista/submit.shtml). CNS: conserved non-coding sequences (conservation cutoffs: identity >= 99% and length >= 100 bp).
